# A signal detection theoretic demonstration of hiring rate asymmetries in competitive academic job markets

**DOI:** 10.1101/061200

**Authors:** Michael Miuccio, Ka-yuet Liu, Hakwan Lau, Megan A. K. Peters

## Abstract

To get a faculty job, graduating doctoral students have to substantially outperform their peers, given the competitive nature of the academic job market. In an ideal, meritocratic world, factors such as prestige of degree-granting university ought not to play a substantial role. However, it has recently been reported that top-ranked universities produced about 2–6 times more faculty than did universities that were ranked lower (Clauset, Arbesman, and Larremore 2015), necessitating un-meritocratic factors: how could students from top-ranked universities be six times more productive than their peers from lower-ranked universities? Here we present a signal detection model to demonstrate that substantially higher rates of faculty production need not require substantially (and unrealistically) higher levels of student productivity. Instead, it is a high hiring threshold due to keen competition that causes small difference in average student productivity between universities to result in manifold differences in placement rates. Under this framework, the previously reported results are compatible with a purely meritocratic system. As a simple proof of concept, we examined the association between university ranking and the impact factors of students publications from a small selected sample of psychology departments in the U.S. The results are in agreement with our theoretical model. Whereas these results do not necessarily mean that the actual faculty hiring market is purely meritocratic, they highlight the difficulty in empirically demonstrating that it is not so.

## Introduction

Is academia a pure meritocracy? If it is not, what makes it deviate from the ideal? Doctoral students now seem to have to substantially outperform their peers in the competitive academic job market to get a faculty position, and the prestige of the degree-granting institution appears to be a crucial factor. A recent study found that top-ranked universities produced about 2–6 times more faculty than did universities that were ranked lower (Clauset et al. 2015). A multitude of mechanisms, including non-meritocratic factors, have been suggested to underlie the differences in placement rates across institutions of varying status, e.g., nepotism, racism, and sexism, and, at the institutional level, hiring network structures and prestige of the programs (Burris 2004; Clauset et al. 2015; Mai, Liu, and González-Bailón 2015; Merritt and Reskin 1997; Misra, Kennelly, and Karides 1999; Wennerås and Wold 1997).

While these previous studies provide good evidence that the academic market may not be a pure-meritocracy, the role of the increasingly high level of competition of academic job markets in generating the observed uneven distribution of jobs as a function of institutional prestige is less well discussed. It is reasonable to expect the qualifications of successful candidates to increase with the level of competition in the job market. That said, one might question whether students from top-ranked universities could outperform their peers by as much as six times in productivity to justify the six-fold difference in placement rates (Clauset et al. 2015). (Certainly, it has been pointed out that the mapping of success and quality (however it is measured) does not have to be linear *even* in a purely meritocratic system. Curvilinear relationships between productivity and return are not unique to the faculty hiring markets, and have been referred as the “superstar” (Rosen 1981) and “winner-take-all” effects (Frank and Cook 1996))

Our contribution to this broader literature is that we use a simple mathematical model of binary decisions to quantify the relationship between the competitiveness of the market and the return rates to productivity. Using faculty hiring as an example, we show that a signal detection theoretic argument is consistent with the large discrepancies in faculty production while supposing largely similar rates of productivity between top-ranked and lower-ranked universities.

Signal detection theory (Green and Swets 1966; Macmillan and Creelman 2004) was developed in World War II to study the detection of information bearing signal in radar when there is noise in the system. Psychologists have also picked up it as a mathematical model to quantify how humans make binary decisions when there is uncertainty. One of the key insights, regardless of the system or observer, is that the criterion that is being used to make the decision is as important as the information in the signal itself.

According to signal detection theory, a decision maker uses a criterion to set a threshold, classifying evidence values above that threshold as belonging to one category, and values that fall below it as belonging to the other category. Setting a high criterion can result in large differences in the *proportion* of evidence that surpasses the threshold for two seemingly similar distributions. Figure 1 shows example distributions representing the height of males (red) and females (blue), with males on average 5 inches taller than females (standard deviations are arbitrarily set to 5 inches for both distributions). For the purpose of explanation, let’s suppose that being 73 inches (6’1”) tall or more qualifies you to be considered a tall person. A criterion is then placed at 73 inches to determine the proportion of males and females who qualify as being tall. With this high criterion set, it becomes clear that the *proportion* of males who qualify as being tall (the area above the criterion under the red curve) is over ten times larger than the *proportion* of females who qualify (the area above the criterion under the blue curve), despite the fact that the distributions’ means differ only by a few inches. A lower criterion (e.g. you’re tall if you are 60 inches tall or more) would not produce such an extreme asymmetric effect, with the proportion of ‘tall’ males being only 1.6 times the proportion of ‘tall’ females.

**Figure 1.**
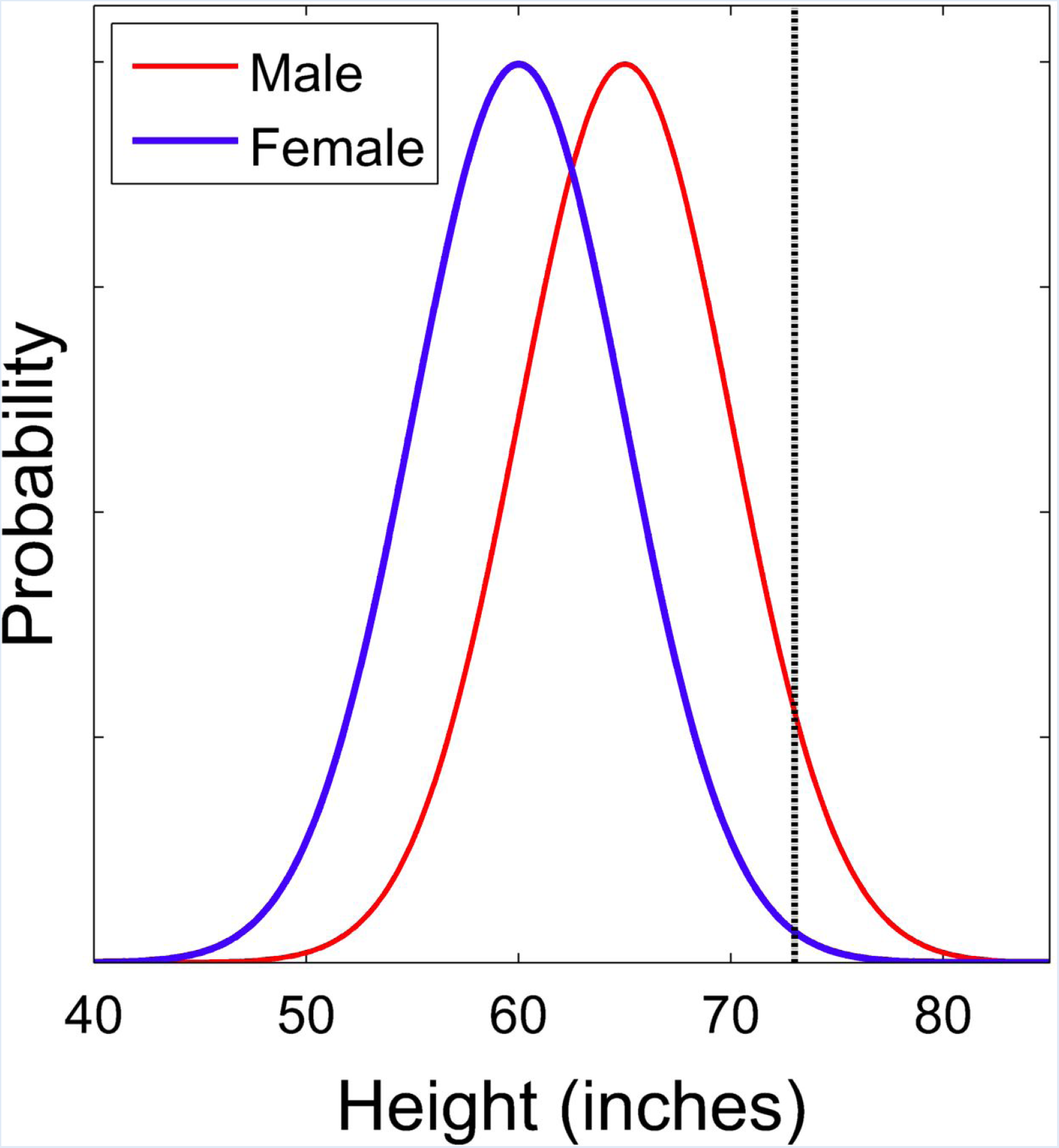
Illustration of the effect of an extreme criterion on categorization according to Signal Detection Theory. Shown is an arbitrary, imaginary example of height distributions for men and women. The distributions differ in mean by only 5 inches, but if the criterion for labeling an individual as “tall” is set to an extreme value, such as 73 inches (6 ft 1 inch), the proportion of males categorized as “tall” (the area above the criterion under the red curve) will be *many* times (in fact, 10.8 times) more than the proportion of females categorized as “tall” (the area above the criterion under the blue curve).

We formalized this theoretical argument as a signal detection model, utilizing the distributions of meritocratic measures of productivity that are likely to exist in true doctoral student populations. We then demonstrated the validity of the theoretical argument by collecting a small sample of graduate students from selected psychology departments in the United States as a proof-of-concept exercise.

Supporting the conditions of the signal detection theoretic model, we found that the productivity of graduate students in the high- and low-tier universities in our sample is actually very similar. However, also in support of the model, faculty production appeared disproportionately skewed toward higher-tier schools. While we cannot rule out the roles of other factors in faculty hiring decisions, such as institutional prestige or social networks as has been previously suggested, the result is also consistent with a high set criterion for faculty hiring. In other words, whereas the model -- and these sample data -- do not necessarily mean that the actual faculty hiring market is purely meritocratic, they highlight the important role of an extreme hiring criterion due to a keenly competitive environment. They also reveal the difficulty in empirically showing that the faculty job market is demonstrably “corrupted” by un-meritocratic factors.

## Signal Detection Theoretic Model

Our signal detection theoretic model proceeds much as the above example of height differences between men and women. In signal detection theory, the *samples* of data available to a decision maker are assumed to have been drawn from distributions with known shape and parameters (Green and Swets 1966; Macmillan and Creelman 2004). For the purposes of examining faculty hiring rates, we assume that these samples represent some form of meritocratic productivity scores given to individual students, such that students from higher ranking institutions constitute a distribution of Higher Tier productivity scores and those from lower ranking institutions constitute a distribution of Lower Tier productivity scores. In our model, we will refer to these productivity score samples for individuals as *x*, i.e. samples of the random variables *X_Higher tier_* and *X_Lower tier_*. Because meritocratic probability scores cannot be negative and are positively skewed, we can assume the known shape of these random variables to be exponential, such that for all students *f(x) ∝ λ e^−λx^*. The proportionality occurs because the distribution is normalized such that they constitute probability density functions. (See Proof-Of-Concept Example, below, for validation of these assumptions.)

Following the height differences example above, if a student has a productivity score *x* that falls above an acceptable value -- i.e., a hiring criterion -- then he or she will be hired as a faculty member. If a student’s score *x* does not exceed the hiring criterion, he or she will not be hired. (Of course, the hiring criterion is a soft criterion, in that students who do not meet or exceed an arbitrary cutoff may still be hired as faculty. However, to simplify the theoretical exercise, we assume the criterion to be a hard boundary.)

To determine the magnitude of this hiring criterion, we first must determine the probability of being hired as faculty. A simplistic formula for this probability of being hired, *p(hire)*, would be to compare the average number of students who graduate from each per year to the average number of faculty hires made at those same universities per year. Thus, we can define for a given year

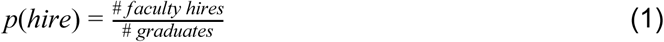

The hiring criterion is defined as the meritocratic productivity score at which the area above this criterion under the distribution of *all* students’ productivity scores regardless of university ranking, *f(x)*, matches *p(hire).* Thus, mathematically the criterion *c* is defined as 

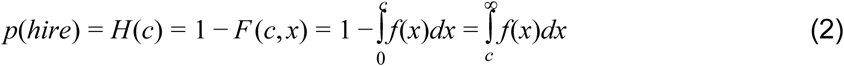
leading to 

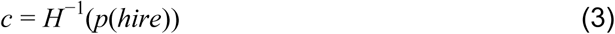
where the integral initially is taken from 0 to *c* because an exponential function is undefined at *x* < 0.

To evaluate how the criterion will differentially affect the conditional probability of being hired as faculty depending on degree-granting institution, *p(hire|university tier)*, we examine the proportion of each conditional probability density function for each university ranking tier that falls above the criterion *c*. Conditioned by the rank of the student’s training institution, the distribution of meritocratic productivity scores is *f(x|tier) ∝ λ_tier_e^−λ_tier_x^*, and thus the conditional probability of being hired is defined as

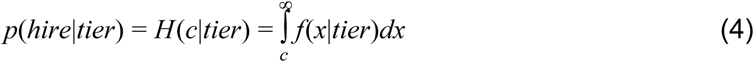

It is assumed that *p(hire)* is quite low due to the competitiveness of the academic job market, with under 10% of graduates with doctoral degrees being hired as faculty members (van Dijk, Manor, and Carey 2014) (see also our Proof-of-Concept Example for validation). At this low *p(hire)*the criterion for being hired will be quite high, as a small proportion of the distribution of all meritocratic productivity scores ought to fall above the criterion. Under a high criterion, modest differences in students’ productivity between universities tiers (quantified as differences in parameter values λ_*tier*_) can lead to radically imbalanced areas under Higher and Lower tier functions above the hiring criterion, *p*(*hire|tier*) -- just as with modest mean height differences between males and females in the example in the Introduction. And, as with the height differences example, a less extreme criterion leads to less asymmetry in hiring rates.

Thus, the signal detection theoretic model predicts the following: (a) students from Higher and Lower tier universities will have similar levels of productivity; (b) under an extremely competitive academic job market with a very high hiring criterion, small differences in student productivity across university tier will lead to manifold differences in probability of being hired; and (c) under a less competitive job market, when the hiring criterion is less extreme, these asymmetries will be reduced.

## Proof-of-Concept Example

### Data collection

Rather than demonstrate the predictions of the signal detection theoretic model with arbitrary simulated data, we elected to demonstrate a more illustrative proof of concept by using a realistic sample of Psychology students’ productivity across universities of different rankings on the *U.S. News & World Report* (2016) list. This discipline was chosen because Psychology has little ‘leakage’, i.e. there are minimal differences between the department discipline an individual graduates from and the department discipline in which he or she is ultimately hired. Psychology also has the additional benefit of an easily-defined (albeit simplified) objective meritocratic measure based on the impact factors of peer-reviewed publications for each individual.

From universities listed on the National Universities Rankings from the *U.S. World and News Report* (2016), we collected three samples of individuals in their Psychology departments depending on university rank — rank 1–10, rank 11–20, and rank 21–100 — through a combination of online search of student directories and direct contact with departments. Overall, we collected data on 1848 individuals from 26 institutions.

### Defining a meritocratic measure

For a meritocratic productivity score, we defined the simple metric of Impact Factor Sum (IFS) for each individual in this sample as the sum of the impact factors for every publication that individual had authored regardless of authorship order, as indexed in Google Scholar. This method was used as a means to equitably search all students’ publications because not all students post their CVs, and data downloaded from large archives (e.g., PubMed) would by definition exclude individuals with no publications and journals not indexed by that engine. We also wanted to quantify productivity for all current students, not graduates.

Although other more complete and complex indices are available -- e.g., h-index (Hirsch 2005, 2007) or predictions based on machine learning techniques (Acuna, Allesina, and Kording 2012) -- we elected to use this IFS metric due to its simplicity and close relationship with more complex metrics. Specifically, it has been shown that the perceived quality (i.e., impact factor) of a publication is given more weight in the faculty hiring process than its actual quality (i.e., its citation rate), and that the two most important factors in predicting faculty hiring are impact factor of publications and number of publications (van Dijk et al. 2014); we therefore combined these factors into a single IFS. Although this simplified IFS metric does not cover all possible facets of meritocratic success, we remind the reader that these data are intended to provide a proof of concept illustration of our signal detection theoretic model.

### Defining university tiers

IFS for each individual in our entire sample ranged from 0 (no publications) to 304.637 (many first-, middle-, and last-authorship papers in high-impact journals) (μ = 7.530, σ = 19.403). These were collected across three groups corresponding to the tier of the university from which an individual had received his or her doctorate: rank 1–10 (n = 607, range: 0–304.637, μ = 11.655, σ = 26.929), rank 11–20 (n = 512, range: 0-96.17, μ = 5.312, σ = 12.471), and rank 21-100 (n = 729, range: 0-150.87, μ = 5.653, σ = 14.872).

Because these samples are not normally distributed, we used nonparametric tests to compare them. We tested for differences among the three samples (rank 1–10, 11–20, and 21–100) with (a) Wilcoxon Rank-Sum (Mann-Whitney U) tests, which are nonparametric tests that do not rely on assumptions of normality (Wilcoxon, 1945), and (b) Kolmogorov-Smirnov nonparametric tests, which test for differences between probability distributions and are sensitive to both mean and distribution shape (Chakravarti, Laha, and Roy 1967). Wilcoxon Rank-Sum (Mann-Whitney tests revealed significant differences in IFS between universities with rank 1–10 and those with rank 11–20 and 21–100, but no differences between universities with rank 11–20 and 21–100 (Table 1, top three rows). Kolmorogov-Smirnov tests revealed an identical pattern (Table 1, top three rows).

**Table 1.** Results of nonparametric comparisons of all university tier groups. All individual groups’ pairwise comparisons are significant (denoted with *) except for rank 11–20 vs. 21–100 (top three rows). We therefore collapsed across the two similar lower groups to create a single pair of groups (bottom row). This pair of groups -- Higher vs. Lower tier -- was used in all further analyses.

|  |  |  |  | Wilcoxon Rank Sum / Mann-Whitney U |  | Kolmogorov-Smirnov |  |
| --- | --- | --- | --- | --- | --- | --- | --- |
|  | Rank pairing | n<sub>1</sub> | n<sub>2</sub> | U | p | D | p |
| **Individual groups** | 1–10 vs. 11–20 | 607 | 512 | 3.596e5 | < .001* | 0.141 | < .001* |
|  | 1–10 vs. 21–100 | 607 | 729 | 4.355e5 | < .001* | 0.156 | < .001* |
|  | 11–20 vs. 21–100 | 512 | 729 | 4.507e5 | .706 | 0.038 | .760 |
| **Merged groups** | Higher (1–10) vs. Lower (below 10) | 607 | 1241 | 6.105e5 | < .001* | 0.149 | < .001* |

Because these tests revealed indistinguishable IFS distributions for universities with ranks 11-20 and 21-100, we collapsed across the two lower-ranking groups to create a Higher tier sample (n = 607, rank 1–10) and a Lower tier sample (n = 1241, rank below 10). The range of IFS scores for the Higher tier was therefore 0–304.637 (μ = 11.655, σ = 26.929), and for the Lower tier was 0–150.87 (μ = 5.512, σ = 13.928). These two tiers are significantly different from each other (Table 1, bottom row).

### Comparing university tiers on meritocratic measures

We evaluated the similarity between the remaining Higher tier (rank 1–10) and Lower tier (rank below 10) groups using Receiver Operating Characteristic (ROC) analysis, which plots the *hit rate* versus *false alarm rate* at varying criterion values (Green and Swets 1966; Macmillan and Creelman 2004): for all possible criterion values, IFS values are defined as *hits* if they are correctly classified as belonging to the Higher tier group, and *false alarms* if they are classified as belonging to the Higher tier group but actually came from the Lower tier group. At each possible criterion value the *hit rate* and *false alarm rate* are calculated, which are then plotted against each other to form the Receiver Operating Characteristic (ROC) curve. The area under this ROC curve (AUC) is a measure of the similarity between the Higher and Lower tier IFS scores, as it provides a normalized metric of separability of distributions: AUC = 0.5 indicates distributions are identical, and AUC = 1 indicates distributions are completely separable.

Despite significant differences between the Higher and Lower tiers at the statistical level, the samples appear visually similar (Figure 2a), and ROC analysis (Green and Swets 1966; Macmillan and Creelman 2004) showed that the normalized magnitude of this difference was indeed quite slight, at AUC = 0.566 (Figure 2b). The interpretation of this AUC value is that if a random person is picked from either cohort and you are asked to guess the cohort to which the person belongs based on his or her IFS score, your likelihood of being correct would be just 6.6% above chance (chance = 50%). This similarity between the two cohorts is also reflected by the median score for both distributions: both Higher and Lower tier groups have a median IFS of 0. So the distributions’ differences are highlighted primarily in the extreme end of their upper tails, mimicking the 5-inch height difference between males and females in the example in the Introduction and demonstrating the first model prediction. All IFS data are available in Supplementary Table 1 (available online).

**Figure 2.**
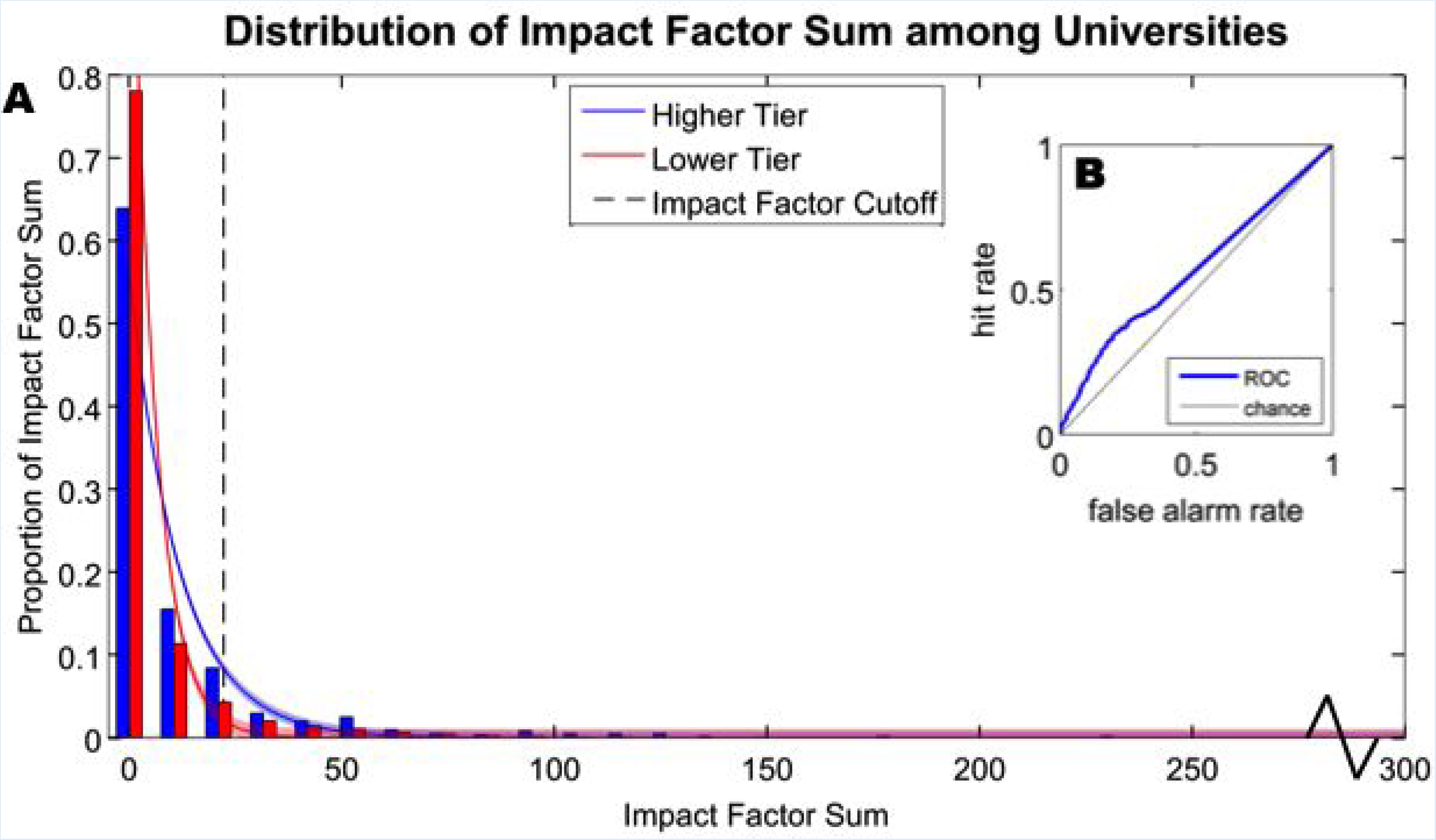
Distributions of Impact Factor Sum (IFS) across university tier are very similar. Panel (a) shows that productivity (graduate students’ IFSs) of the different university tiers (Higher vs. Lower) is quite similar while the criterion for getting a faculty position is, as expected, fairly high. The difference between the two IFS distributions for Higher and Lower tiers is minimal, as shown by the ROC curve in (b): the area under the curve (AUC), representing discriminability between Higher and Lower tier universities, is 0.566. This is almost at chance (chance AUC = 0.50), showing that the distributions are nearly equivalent, mimicking the 5-inch height difference between males and females in our simple example in the Introduction. To calculate the location of the IFS criterion to be hired as faculty (i.e., IFS Cutoff), we fitted an exponential function to the overall distribution of IFS, which represents productivity of all graduate students regardless of university (fitted curve not shown). The IFS Cutoff was calculated to be 22.20 in accordance with the reality of faculty production. The percentage of the area under the two curves representing the Higher and Lower tiers that falls above this IFS Cutoff, *i.e.,* the probability of a graduate student getting a faculty position after graduating from *any* university, is about 5%.

### Estimating the probability of being hired as faculty

To define the criterion in IFS space for being hired as faculty regardless of graduate university, as described in the Model section above, we estimated the probability of being hired as faculty across a large number of universities. To do this, we collected a fourth sample from the Psychology departments across all *U.S. News and World Report* ranks from 1–50, to compare the average number of students who graduate from each per year to the average number of faculty hires made at those same universities per year.

In this sample of graduates and hires from Psychology departments at 22 institutions, we found that an average of 682.6 students graduated per year, and an average of 35.5 individuals were hired as faculty at those same institutions per year. By Equation 1, this leads to *p(hire)* = 0.0520, meaning that approximately 5% of all graduates with doctorates in Psychology are hired as faculty in any given year. This result is in line with previous reports of faculty hiring rates of about 6.2% (van Dijk et al. 2014).

### Setting the criterion for being hired as faculty

To calculate a criterion for being hired within the overall IFS distribution, we collapsed all IFS for all individuals in the Higher and Lower tier groups regardless of university tier. We then used bootstrapping to ensure that our demonstration of the model’s predictions was not overly sensitive to our particular sample. On each bootstrap loop, a random sample of 1000 IFS data points (with replacement) was drawn from this overall IFS distribution. To each sample of 1000, we fitted an exponential function of the form *f(x) ∝ λ e^−λx^* to all data regardless of university tier, which we then normalized so that it would constitute a probability density function over the range of IFS in our sample (see Signal Detection Theoretic Model, above). We then calculated the criterion *c*, or IFS Cutoff, according to Equations 2 and 3. This process was repeated 1000 times for a total of 10,000,000 samples, leading to 1000 estimates of *c.* We found mean *c* = 22.20 (median = 22.20, σ = 1.74) (Figure 2), meaning that, if we live in a meritocracy, any given individual should aim to have a total IFS equal to or exceeding 22.20 if he or she hopes to be hired as a faculty member.

### Extremely unequal probability of being hired as a function of university tier

Despite the visual similarity between the Lower and Higher tier distributions of IFS (Figure 2a), closer inspection of the tail ends of the distributions, above the criterion *c*, reveals important differences. Figure 3a displays the mean of *f(x|tier)* for both tiers over all loops of the bootstrapping analysis zoomed in on the region of the IFS Cutoff criterion *c*, with SEM across bootstrap loops represented by the shaded regions. The Higher tier IFS scores display a strikingly large advantage over the Lower tier IFS scores at the location of the criterion.

**Figure 3.**
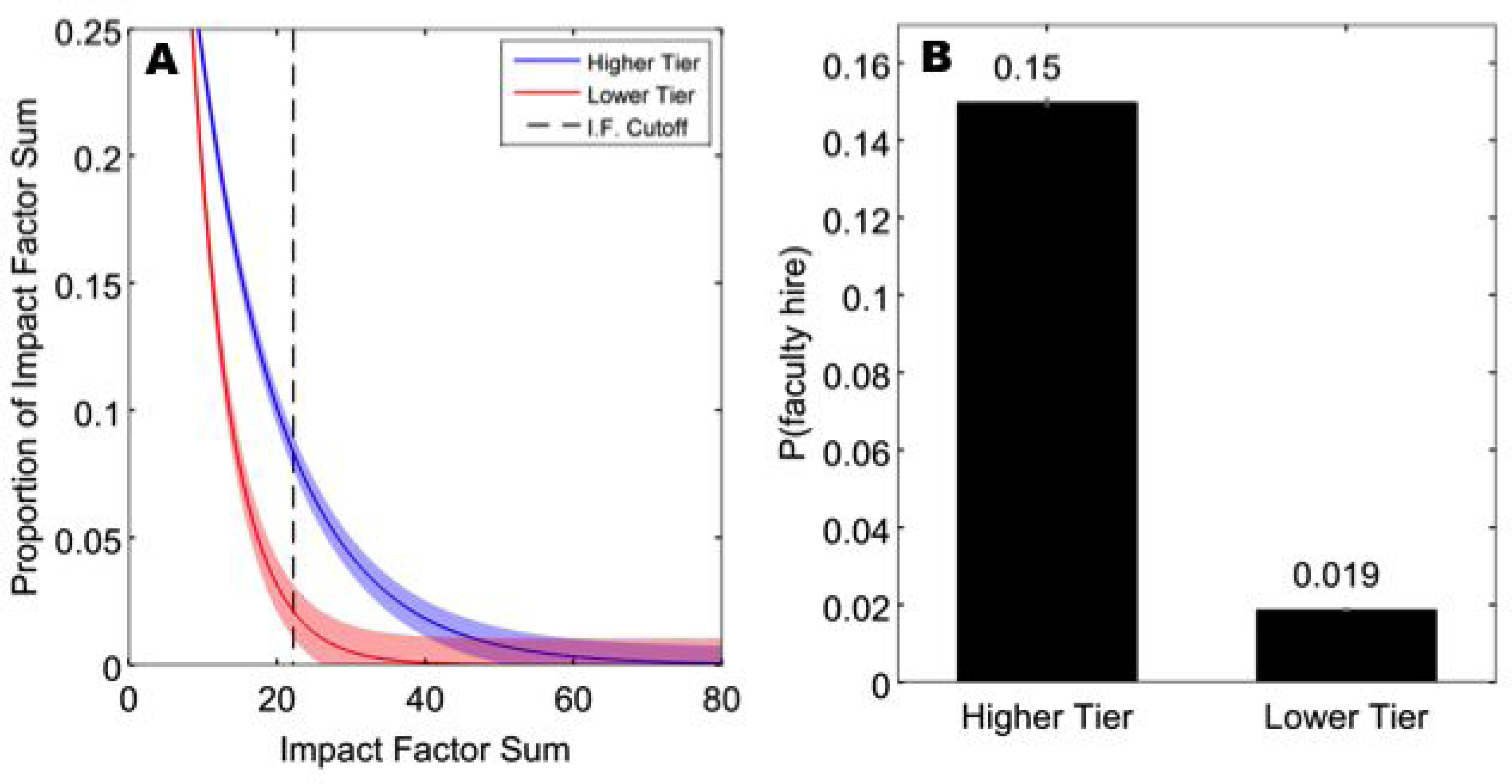
Zoomed-in view of IFS distributions and probability of faculty hire conditioned on university tier, *p(hire|tier)*. Panel (a) shows the portion of the graph from Figure 2a nearest the criterion. At this zoom level it is clear that despite their overall similarity, the distributions are quite different at the relatively extreme value of the IFS criterion. Shaded regions indicate the standard error of the mean (SEM) across bootstrapping loops (see Methods) at each IFS value. Panel (b) shows the average probability of being hired as faculty conditioned on having graduated from a Lower or Higher tier university, or mean *p(hire|tier)* across bootstrapped samples (see Methods). According to our sample, the mean probability of being hired after graduating from a Higher tier university is 14.98%, or about eight times the mean probability of being hired as faculty after graduating from a Lower tier university (1.88%). Error bars represent the SEM across all bootstrapping loops.

To evaluate how the high criterion might lead to potentially exaggerated differences if we conditioned on university tier, i.e. *p(hire|tier)*, we again used bootstrapping to ensure that our results were not overly sensitive to our particular sample. As before, on each bootstrap loop, a random sample of 1000 IFS data points (with replacement) was drawn from each of the Higher and Lower tiers, respectively, and an exponential function fit to the sample of the form *f(x)∝ λ_tier_e^−λ_tier_x^*. Following normalization such that the fitted functions constituted probability density functions, we used Equation 4 on each bootstrap loop for each tier to calculate the probability of being hired conditioned on university tier. This process was repeated 1000 times for a total of 20,000,000 samples (10,000,000 from each of the Higher and Lower tiers), leading to 1000 estimates of *p(hire|Higher tier)* and 1000 estimates of *p(hire|Lower tier)*.

Figure 3b displays the mean and standard error of *p(hire|tier)* for the Higher and Lower tier universities across all loops of the bootstrapping analysis. Graduates of Higher tier universities are significantly more likely to be hired as faculty (t(999) = 157.627, p < .001), and by nearly a factor of eight: you are almost eight times as likely to be hired as faculty if you receive your doctorate in Psychology from a top-10 university than if you attended any university of lower rank, based purely on the simplified, meritocratic metric of IFS. This occurs despite the high degree of similarity in the IFS distributions for Higher and Lower tier universities (AUC = 0.566), as a result of small but significant differences in the probability densities of these distributions at and above the high hiring criterion. This demonstrates the second model prediction.

### Asymmetry in hiring rates is a direct consequence of an extreme criterion

An important lesson from our signal detection theoretic model is that the extreme criterion (i.e., low probability of being hired as faculty) set by a keenly competitive faculty job market is largely responsible for the large asymmetry in faculty hiring rates. The effect of the extreme criterion is clear in our proof-of-concept example dataset, the criterion for being hired at IFS ≈ 22 reflects the externally valid hiring rate of ∼5–6% (van Dijk et al. 2014), and leads to hiring asymmetry by a factor of nearly eight between Higher and Lower tier universities.

If the hiring climate were less competitive, with a less extreme hiring criterion along any meritocratic dimension, this asymmetry would dwindle with increasing values for *p(hire)* and eventually disappear (Figure 4). To illustrate this consequence of the signal detection theoretic model, we calculated the hiring rate asymmetry between Higher and Lower tier universities (Equations 3 & 4) as a function of increasing *p(hire)*, i.e. decreasing competitiveness of the academic job market. As above, we also used bootstrapping analysis with 1000 samples of IFS scores from the overall distribution. Although the mean *p(hire)* ratio between Higher and Lower tier universities starts high, as would be expected in the current competitive hiring climate, it dwindles rapidly with decreasing competitiveness, asymptoting as *p(hire)* approaches 1. This demonstrates the third model prediction. Thus, the *appearance* of a non-meritocratic system is in fact perpetuated in large part by the extreme difficulty of attaining a faculty position.

**Figure 4.**
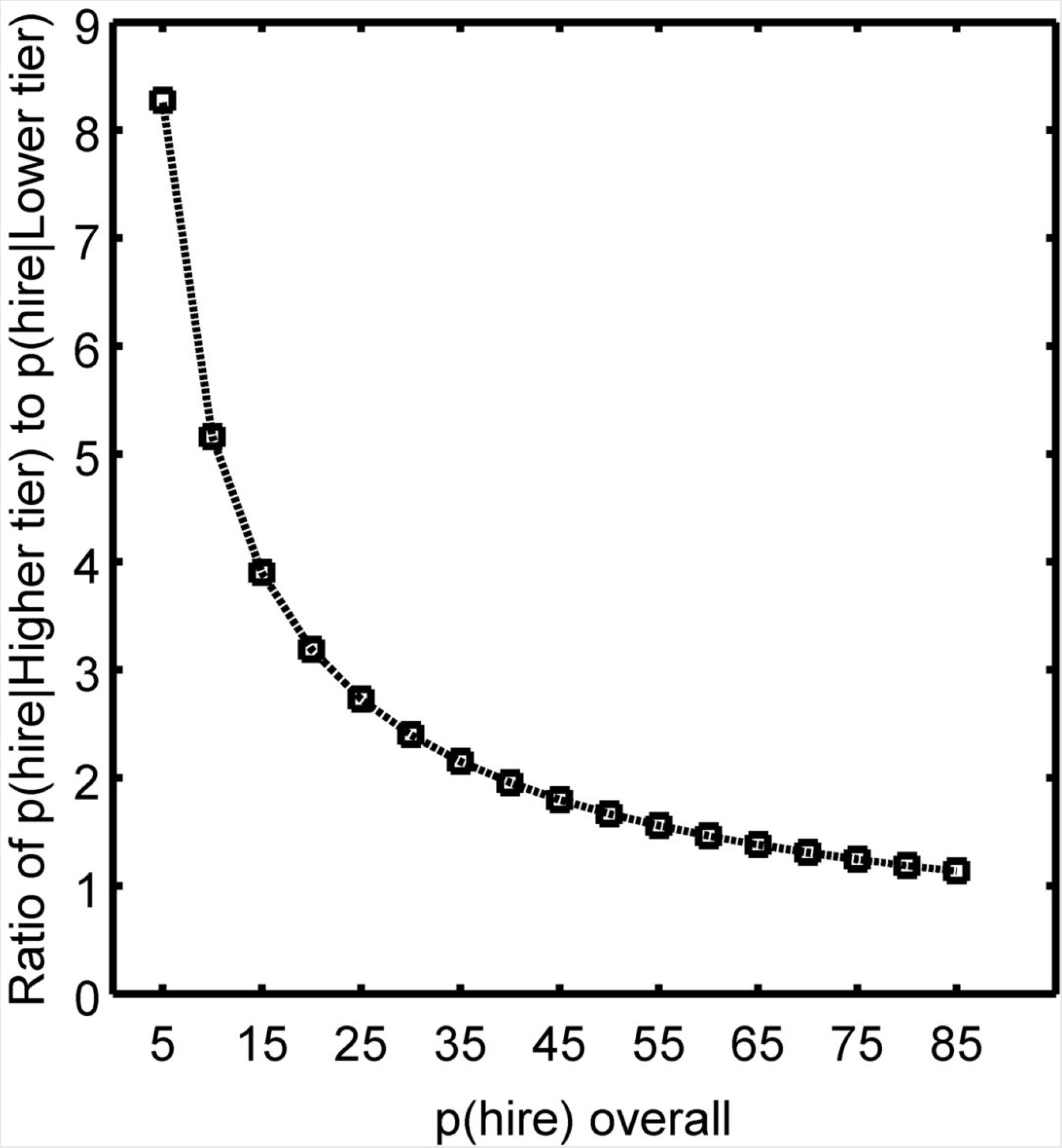
Less extreme hiring criterion values lead to less pronounced hiring asymmetry between university tiers. By shifting the criterion to more liberal values — from *p(hire)* = 0.05 to *p(hire)* = 0.85 — we show that the hiring asymmetry is reduced and ultimately disappears almost entirely. The appearance of a non-meritocratic system is thus perpetuated by the severe competitiveness of the current hiring climate.

### Effect of focusing on more advanced students

To demonstrate that the validity of our signal detection theoretic model is not dependent on the idiosyncrasies of any particular sample, we repeated the above-described analyses excluding individuals with IFS = 0 from both Higher tier and Lower tier samples (n_Higher_= 272, n_Lower_= 448). This can be thought of as selecting primarily the more advanced students while removing first- and second-year students who have not published yet, to alleviate concerns that the proof-of-concept results reflect the relatively large proportion of students who had no publications at the time of our data sample collection.

Analyses on this subsample of individuals shows no change in overall findings. Firstly, higher tier universities still stand out from universities with lower rankings in terms of student productivity (Table 2), justifying the collapsing of the lower two university tiers. Further, just as in the main analysis, despite a higher IFS Cutoff (criterion) at 57.05, resulting from the shift of probability density towards higher IFS values, the observed similarity between Higher and Lower tier distributions is maintained (albeit a little lower, ROC = 0.6358), and resultant ratios of Higher to Lower tier faculty hiring rates are similarly starkly asymmetric: mean *p(hire|Higher tier)* = .112, mean *p(hire|Lower tier)* = .024. This result demonstrates that while our sample may be small and IFS Score may not capture all possible meritocratic elements, the proof of concept of our signal detection theoretic argument does not depend on any specific sample. See Supplemental Material for further discussion.

**Table 2.** Results of nonparametric comparisons of Higher and Lower tier groups after individuals with IFS = 0 have been removed.

|  |  |  |  | **Wilcoxon Rank Sum / Mann-Whitney U** |  | **Kolmorov-Smirnov** |  |
| --- | --- | --- | --- | --- | --- | --- | --- |
|  | **Rank pairing** | **$n_1$** | **$n_2$** | **U** | **p** | **D** | **p** |
| **Individual groups** | 1–10 vs. 11–20 | 272 | 186 | 6.891e5 | < .001* | 0.212 | < .001* |
|  | 1–10 vs. 21–100 | 272 | 262 | 8.281e5 | < .001* | 0.251 | < .001* |
|  | 11–20 vs. 21–100 | 186 | 262 | 5.753e5 | .339 | 0.098 | .239 |
| **Merged groups** | Higher (1–10) vs. Lower (below 10) | 272 | 448 | 1.115e5 | < .001* | 0.233 | < .001* |

## Discussion

Here, we have used a simple signal detection model (Green and Swets 1966; Macmillan and Creelman 2004) to demonstrate how surprisingly extreme asymmetries in faculty hiring rates for graduates of Lower and Higher tier universities need not necessitate non-meritocratic factors in the faculty hiring process. Importantly, the theoretical model does not depend on particularities of distribution shape, distribution parameters, or sample size. Simply put, as long as the faculty job market is competitive -- i.e., a high meritocratic criterion exists because the probability of being hired is low -- small differences in productivity as a function of university rank will be magnified into manifold differences in faculty production rates. While it has been shown that doctoral prestige better predicts faculty placement than productivity (Burris 2004; but see van Dijk et al. 2014; Miller, Glick, and Cardinal 2005), our theoretical argument casts doubt on the supposition that extreme hiring asymmetries *must* imply either vast (and unrealistic) differences in productivity, or a strongly non-meritocratic system.

We believe that unmeasured heterogeneities in the positions or in the candidates are certainly important in the hiring decision. That said, our model provides an alternative explanation to why controlling for publication records usually has only modest effects on the prestige-placement relationship (e.g., Headworth and Freese 2015) More importantly, our study shows that, in principle, such small differences can aggregate into large differences in the placement records across institutions, even in a pure-meritocracy.

That decision-makers likely consider factors beyond Impact Factor Scores in the selection process may explain why the observed differences in placement across institutions are less than what our model predicts. It may also explain the heterogeneity we observed in actual faculty hiring decisions. However, what our modeling exercise demonstrates is the difficulty in quantifying the importance of unobserved factors when the market becomes extremely competitive.

### Proof-of-Concept Example

We used a realistic sample of data to demonstrate the consequences of our signal detection theoretic model, which revealed that true rates of faculty production are highly skewed towards Higher tier universities by a factor of eight even though student productivity between tiers is nearly indistinguishable (AUC = 0.566). The high criterion (IFS Cutoff) in our sample (dictated by an extremely competitive faculty hiring system) is reflective of actual faculty hiring rates both in our sample and as reported by others (5-6%; van Dijk et al. 2014), thus providing a realistic dataset with which to demonstrate the predictions of our signal detection theoretic model. We also showed that if this hiring criterion were not so extreme, asymmetry in faculty hiring rates as a function of university rank would dissipate.

Our example has several limitations. One possible limitation is the means by which we gathered data to calculate IFS based on Google Scholar publication results. We elected to collect publication data via Google Scholar because (a) individuals do not have control over what appears in the search engine (unlike their appearance in NeuroTree or uploaded CVs) and (b) we wanted a metric by which to quantify all students’ productivity as a function of university tier including students who had published nothing at all. By searching for individual students’ names (collected from student rosters) on Google Scholar we were able to calculate all students’ IFS scores for all papers they had published, if any. This helped to keep our sampling method equivalent among all the universities we sampled from.

Another possible concern is that IFS score metric itself may not capture all relevant aspects of student productivity. We used IFS despite its possible over-simplification because it has been shown to be a strong predictor of success in academic job markets (van Dijk et al. 2014). Further, total impact factor score has been shown to be more predictive of fellowship application success than measures based only on first-author publications or number of citations (Wennerås and Wold 1997). Thus, for the purposes of illustrating the theoretical argument, we favored the IFS metric over h-index (Acuna et al. 2012; Hirsch 2005, 2007) because of its simplicity and similarity to previously validated approaches.

The assumption of our simplified example that graduates seeking faculty positions who are not hired do not cumulatively add to the faculty applicant pool from year to year is certainly unrealistic. It is quite likely that at least some of the remaining 95% of graduate students who are not hired would be added to the following years’ new applicants, while some of them would take up a post-doctoral or a non-tenure track position before getting a tenure-track job. Nonetheless, there are reasons to expect a long transition period and the existing hierarchy between tenure-track and non-tenure track positions would result in an even more extreme criterion, leading to even more inequality in job placement since we expect cumulative advantage (Bedeian et al. 2010; DiPrete and Eirich 2006; Headworth and Freese 2015) to exacerbate the differences in IFS between Higher and Lower tier universities during the pre-tenure track years. Even if the post-doctoral job market is not as competitive as the faculty job market and thus has an equalizing effect, we are skeptical that the effect is enough to compensate for the impact of the current high threshold in the faculty job market.

Despite these potential criticism of the methods we used to collect and define our proof-of-concept dataset, the predictions of the signal detection theoretic model do not depend on these choices. The signal detection theoretic model -- demonstrating that a small difference in meritocratic measures across university rank can lead to a manifold difference in hiring rates under an extreme criterion -- holds in any distribution, and so the sample we gathered serves primarily to provide a concrete example of our theoretical argument.

### Conclusions

We have shown that similarly productive cohorts will produce very different rates of faculty hires simply because of a high hiring threshold, by demonstrating the predictions of a theoretical model with a realistic dataset. It should be noted that our results *cannot* definitively speak to whether or not the current system is a in fact pure meritocracy (see e.g., van Dijk et al. 2014 for discussion of the impact of university rank and gender on faculty hiring). However, our demonstration does make clear that a substantial discrepancy in hiring rates as a function of degree-granting university tier *may not*, in fact, *necessitate* factors beyond the meritocratic.

## Acknowledgements

This research is partially supported by the National Institutes of Health (NINDS R01 NS088628-01 to HL). We also thank Nikolaus Kriegeskorte for seminal discussion on internet social media which inspired the study.

